# Physiological and Metabolic Responses of Chickpea to Post-Flowering High Temperatures and Limited Water Availability

**DOI:** 10.1101/2025.08.28.672843

**Authors:** Carla Pinheiro, Eduardo Costa-Camilo, Manzur-E-Mohsina Ferdous, Carina Barcelos, Isabel Duarte, Leonor Guerra-Guimarães, Thomas Roitsch

## Abstract

Post-flowering elevated temperatures are increasingly frequent and strongly affect crop yield and seed nutritional quality. This study examined the impact of elevated post-flowering temperatures (32°C/25°C, day/night) combined with two watering regimes (40% and 10% field capacity) on chickpea genotypes. Elvar and Electra, top-yielding Portuguese genotypes well adapted to southern Portugal’s dry climate were evaluated. Under controlled high-temperature conditions (Phenolab), reproductive cycle was shortened to 30-35 days, compared with 65 days under greenhouse conditions (24°C/18°C). Water use peaked soon after flowering and declined after 19 days, regardless of genotype or watering regime, suggesting a physiological limitation on water use. Elevated temperature strongly reduced seed number and weight while altering composition. Nutrient density improved, with higher protein and mineral levels (P, Mg, S, Mo, Fe, Zn) and lower starch content, highlighting a consistent protein–starch trade-off unaffected by water availability. While growth environment largely determined composition, enzymatic activity patterns revealed genotype-specific differences in carbon metabolism (e.g. sucrose synthase, cell wall invertase, aldolase and phosphoglucose isomerase). Overall, these findings suggest genotype-specific responses driven by carbon metabolism and emphasize integrating metabolic, physiological and composition traits in breeding for yield stability and nutritional value.

**Highlight:** Post-flowering temperature dictates reproductive duration and seed composition, enhancing nutritional density but lowering yield. Protein-starch trade-off was consistent across genotypes, while enzymatic activity profiles enabled genotype discrimination under contrasting environments.

## Introduction

Edible grain legumes are a key ingredient of diets worldwide and provide a rich source of protein and essential minerals. They play a critical role in global food security by supporting initiatives to combat undernutrition, micronutrient deficiencies and diet-related diseases. Among them, chickpeas (*Cicer arietinum* L.) stand out for their high protein content, exceptional nutritional benefits, and versatility in global cuisines. Recognized for having one of the best nutritional qualities among seed legumes it is often referred as functional food (Frimpong et al. 2009; Ribeiro et al. 2017; Wang et al 2021; Ajay et al. 2024; Kumar et al. 2025).

Similarly to other crops, chickpea performance is influenced by soil water availability and high temperatures, particularly during pod set and seed filling (e.g. in Krishnamurthy et al., 2010; in Pang et al., 2017, Sita et al. 2017). In addition, compounded physiological damage due to the co-occurrence of high temperatures and water availability are reported (Awasthi et al. 2014, 2017; Benali et al. 2023), underscoring the necessity of studying such interactions. Well adapted to the climate and agronomic conditions of the Mediterranean basin, chickpeas can exhibit drought and heat tolerance in some phenological stages (Devasirvatham et al., 2015). Nevertheless, water availability remains a major constraint on chickpea grain yield, with reported losses ranging from 36% to 42% (Saxena et al., 1993; Khodadadi, 2013). When combined with elevated temperatures, seed filling is further impaired, leading to even larger yield reductions (Kaushal et al., 2013; Devasirvatham et al., 2015; Devi et al., 2023; Benali et al., 2023). The increasing frequency of high temperatures under limited water-availability during the reproductive phase, driven by climate change, is expected to exacerbate these challenges. A typical response to post-flowering high temperatures is a shift in the duration of phenological phases, resulting in decreased seed set and accelerated senescence (Devasirvatham et al., 2015; Maity et al. 2023), being advocated to adjust sowing dates to minimize stress exposure, as well as the breeding of crop varieties with enhanced tolerance to environmental stresses. In contrast with yield loss, post-flowering high temperature impacts on seed quality are far less documented; to the best of our knowledge, only Thenveetti et al. (2024) reported such effects on soybean. Driven by altered water availability and altered enzymatic activities, changes in source-sink dynamics are anticipated to impact seed nutritional composition and quality by affecting processes such as CO₂ absorption, photosynthesis and sucrose translocation, as well as the absorption and transport of inorganic minerals as nitrogen, iron or zinc (Griffiths et al. 2016; Sehgal et al. 2018; Morin et al. 2022; Maity et al. 2023). In the context of sink strength, genotype specific differences in the activity of several carbon metabolism enzymes in the seeds, involved in sucrose metabolism and starch accumulation are key players in determining how efficiently the seed can import, convert, and store carbon (Weber et al. 2005; Sita et al. 2017).

Considering sustainable yield and nutritional quality under future climate scenarios as key ideotype targets, genotype-specific assessments are crucial for advancing our understanding and guiding breeding programs aimed at enhancing crop resilience and performance. The characterization of current elite chickpea genotypes under post-flowering high temperatures and limited water availability is particularly important for developing strategies to optimize the performance of grain legumes, especially with respect to seed sink strength and reserve allocation. In this context, seed biochemical analysis offers valuable insights into the nutritional consequences of environmental stress, thereby supporting efforts to preserve both yield and quality under increasingly challenging climatic conditions.

## Material and Methods

### Plant material

Two chickpea genotypes of the kabuli type currently included in the Portuguese national catalogue of varieties were used. Elvar, the 1^st^ cultivar developed and available since 1993, is adapted to dry conditions and has a high production potential. Electra was included in the catalogue in 2020 and was selected due to its production potential and large seed size.

Seeds obtained under field conditions at Elvas during the 2019-2020 growing season were used as sowing material for all the plant growing experiments (see details below).

### Field assays

Field assays were performed at “Herdade da Comenda” (part of the Portuguese National Plant Breeding Station in Elvas, Portugal; 38°53’33.4”N; 7°03’15.3”W), under a typical Mediterranean environment and with common wheat (*Triticum aestivum* L.) as a pre-crop. A brief characterization of climate, phenological dates and yield are provided as Supplemental Table S1.

Per genotype, one hectare was sowed each year (7 December in 2020; 8 January in 2021), at a density 25 viable seeds per m^2^ with a 50 cm distance between rows. After sowing, herbicide Pendimetaline (active substance) in the order of 3.0 l ha^−1^, was sprayed. Seeds were harvest at the full maturity stage.

### Controlled conditions assays

At Taastrup field station, greenhouse (GH) and an automated phenotyping platform (Phenolab) were used. In the GH, seeds were sown in Phenolab compatible pots [11×11×11 cm, ca. 1.5L soil volume], filled with subsoil from the Taastrup field station and fertilized with 100 kg N per ha, 28 kg P₂O₅ per ha, and 78 kg K₂O per ha. A total of 65 pots per genotype were used and after emergence, the three seeds sowed per pot were trimmed and a single germinated seedling was left per pot. From 2021.05.31 (sowing) to 2021.06.22, plants were maintained fully irrigated under a 16h/8h photoperiod and 24 °C/18 °C (light/dark cycle). To accelerate flowering, the photoperiod was adjusted to 20h/4h (light/dark) until 2021.07.06, when 90% of the plants had at least one flower. At this point, a subset of 25 plants per genotype was kept in the greenhouse (GH) and 40 plants of each genotype were transferred to the Phenolab for the stress treatments.

At the GH, plants were kept under a 16h/8h photoperiod and 24 °C/18 °C (light/dark cycle), with watering adjusted to 40% field capacity. The growing conditions are available as Supplemental Table S2. Irrigation was fully stopped at the end of the Phenolab assay (2021.08.13) to force growth cycle completion. All plants were completely dry at 2021.09.10, and yellow pods with rattle seeds (harvest maturity) were harvested and considered for further analysis.

At the Phenolab, 2021.07.08 was defined as d01, and following acclimation periods, plants grown at 32 °C/25 °C under mild drought (40% field capacity) or with minimal irrigation (10% field capacity) to mimic in-depth water uptake by roots when plants are grown in soil. The growing conditions are available as Supplemental Table S3. Two acclimation periods were observed to gradually adjust watering and temperature (Figure 1). From d01 to d03 (2021.07.08 to 2021.07.10), the watering regime was adjusted to 40% of the field capacity for 20 plants per genotype. For the other 20 plants, the watering regime was stepwise decreased to achieve 10% field capacity at the end of the acclimation period (40%, 20%, 10%). A second acclimation period was introduced to achieve the target temperature gradually avoiding a sudden heat spell ( 10 °C in a single day). The temperature was increased as follows: d04 (2021.07.11): day 24 °C, night 19 °C; d05 (2021.07.12): day 26 °C, night 21 °C; d06 (2021.07.13): day 29 °C, night 23 °C; d07 (2021.07.14): day 32 °C, night 25 °C.

**Figure 1.**
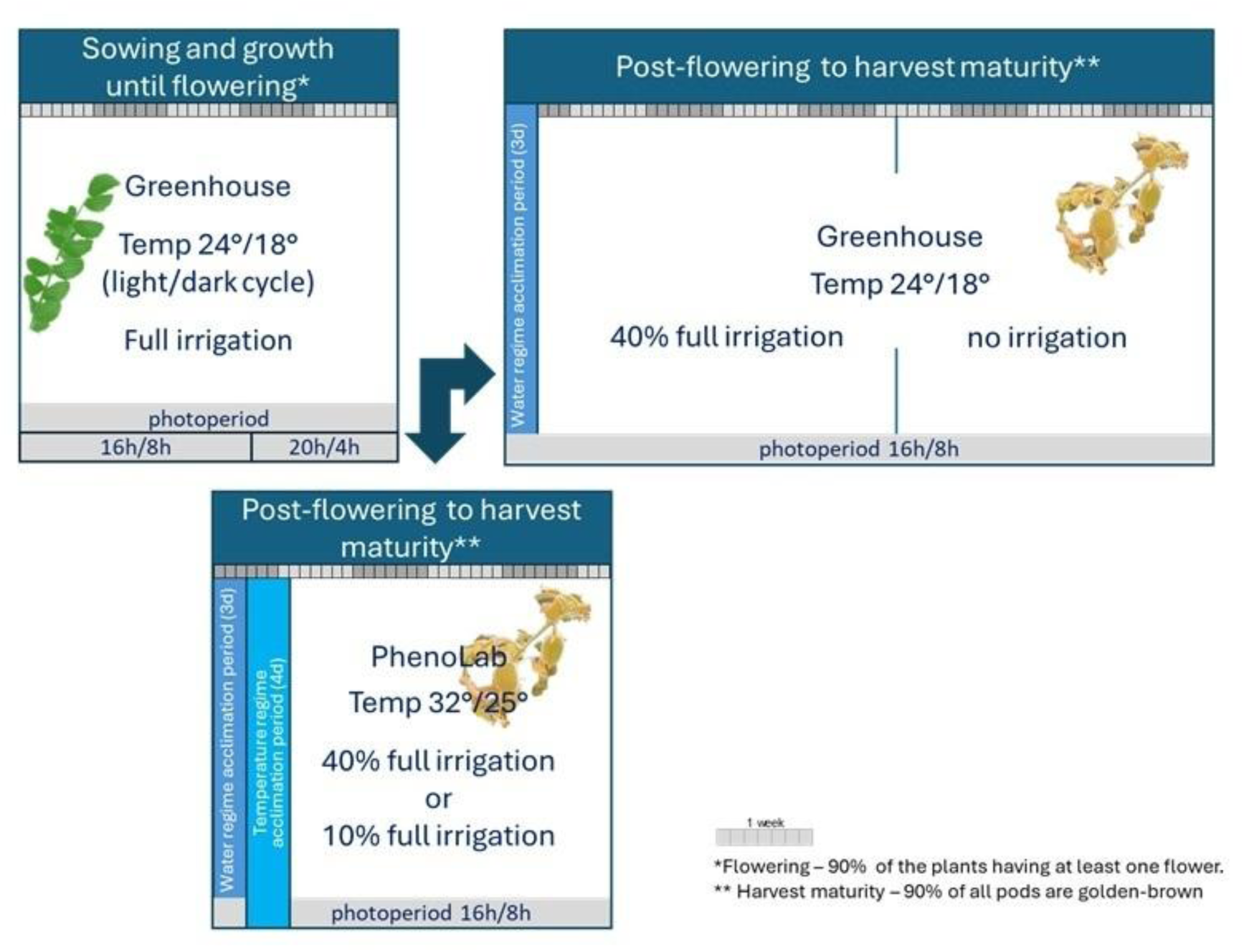
Experimental setup for growing chickpea genotypes Elvar and Electra under controlled conditions at Traastrup, Denmark. The genotypes were cultivated in a greenhouse environment under favourable conditions or subjected to post-flowering high temperatures. The heat stress treatments were combined with two irrigation levels: 40% and 10% of full irrigation.

These conditions (32 °C/25 °C) were maintained from d08 to d35, i.e., until 2021.08.13. Regardless of the watering regime, most plants were completely dry by this date, showing yellow pods and rattling seeds indicating harvest maturity. The Phenolab assay was stopped, and yellow and dry seeds from completely dry plants were collected for further analysis. Growing conditions at the GH and the Phenolab are provided as Supplementary Tables S2 and S3, respectively.

### High-throughput multispectral imaging at the Phenolab

Phenolab standard multispectral imaging procedures were applied. Briefly, images were automatically acquired twice a day by a top-mounted CCD camera with a light-integrating hemisphere setup. The Phenolab imaging unit to make use of ten spectral bands (365, 460, 525, 570, 645, 670, 700, 780, 890 and 970 nm), with a spectral resolution of four megapixels and 2056×2056 pixels. Using the software VideometerLab (version 3.22, Videometer), crop coverage (plant exposed area) was calculated from segmented transformed images and pixel reflectance intensities extracted from the same regions of interest. Thermal imaging was obtained with the FlirA65 thermal camera (FLIR Systems Inc.). Thermographic data extraction was automated by applying masks generated from the segmentation of multispectral images. The multispectral images were geometrically transformed through image registration to align with the thermal images to enable the segmentation of the thermal signal for each plant part acquisition and the calculation of temperature statistics. It needs to be stressed that thermal images were neither normalized nor calibrated, its use being limited.

### Seed analysis

For each genotype and treatment (GxT), seeds were pooled before use for elemental and biochemical analysis. Pooled seeds were ground under liquid nitrogen, aliquoted and kept at −80 °C until use.

### Seed phenotype

A pool of 25 seeds per GxT was photographed under the same lighting conditions. A ColorChecker Passport Photo 2 (X-rite, Pantone) was included in the image for colour calibration and standardization.

#### Germination potential

The germination potential was assessed with a pool of 25-30 seeds per GxT. Germination occurred indoors, at room temperature, under natural light conditions and well-watered conditions, in perforated aluminium trays containing washed and sterilised sand. Seeds were distributed in lines, GxT separated by coloured sticks. Germination success was evaluated 8-10 days after sowing, with all seeds germinating.

#### Elemental analysis

To quantify total nitrogen (N), fat, fiber, ash and individual mineral elements, 15-20g of pooled seeds were used. In brief, total N was performed by Kjeldahl method according to ISO 20483: 2013, using a digestion unit (FOSS Tecator 2520, Denmark), a distillation unit (FOSS Kjeltec 8200, Denmark) and a sequential titration with HCl 0.1N. According to ISO 24557: 2009, moisture content was determined following a 2h oven incubation at 130 °C (Memmert, Germany). According to the ISO 5498: 1981, crude fiber content was estimated using a FOSS Fibertec 8000 system (FOSS, Denmark). Crude fat was estimated according to the AOAC official method 2003.05, following a Soxhlet diethyl ether extraction under reflux (Soxtec system 1043, FOSS Tecator, Denmark). Ash was estimated according to ISO 2171: 2023 by incubating samples in a Nabertherm LT15/11/B510 furnace (Germany) at 550 °C for 4 h. Individual minerals (K, P, S, Mg, Ca, Fe, Ni, Mn, Cu, Mo, Zn, B) were quantified by inductively coupled plasma optic emission spectroscopy (ICP-OES) following microwave-assisted digestion as described in Di Donato et al. (2021). Briefly, 6 ml of 69 % nitric acid was added to 0.2 mg sample and microwave digestion occurred in a closed digestion vessel. A laboratory microwave oven (Multiwave Go, Anton Paar, Austria), equipped with twelve 50-ml vessels made with PTFE-TFM and operating at a frequency of 2455 MHz with an energy output of 850 W, was used for sample acid decomposition. The temperature was linearly increased to 100°C in 10 min and maintained for 2 min; the temperature was linearly increased up to 150 °C in 10 min and maintained for 10 min. After digestion, samples were allowed to cool down and diluted to 10 ml volume with deionized water for ICP-OES analysis, model Plasma Quant 9100 Elite from Analytik Jena (Germany), equipped with Dual View Plasma (radial/axial) and a double monochromator with a CCD detector, cyclonic nebulized chamber and a concentric nebulizer. The operating conditions of the equipment for analyte determination were followed under the manufacturer’s recommended conditions for a radiofrequency power applied of 1.2 kW, plasma gas flow rate of 12.0 l min^−1^ and nebulizer gas flow rate of 0.5 l min^−1^ and calibration was performed using a series of standard solutions with known concentrations of the elements of interest. The analytical wavelengths (nm) chosen were 249.678 (B), 315.887 (Ca), 327.396 (Cu), 259.940 (Fe), 769.897 (K), 285.213 (Mg), 257.610 (Mn), 202.030 (Mo), 231.604 (Ni), 213.618 (P), 180.672 (S) and 213.856 (Zn).

#### Starch quantification

Starch content was estimated after the water extraction of soluble components (adapted from Damesin and Lelarge, 2003). Briefly, 20-25mg of ground seeds were twice extracted in 0.5 ml H_2_O for 20 min under constant stirring (4 °C). After centrifugation (10000*g*, 5min, 4 °C), supernatants were combined and the resulting pellets were further washed by centrifugation (10000*g*, 5min, 4 °C), twice with 70% ethanol and the last one with H_2_O. As described in Pinheiro et al. (2023), starch was gelified in 0.4 ml H_2_0 containing 0.7% (v/v) α-amilase (Sigma A4582) by boiling for 3 min and autoclaving for 1h at 120 °C. After cooling to room temperature, 0.6 ml of sodium acetate buffer (160 mM, pH 4.8) containing 0.28% (w/v) amyloglucosidase (Sigma A7420) was added and samples incubated 2h at 60 °C under constant agitation. After centrifugation (10000*g*, 5min), the supernatants were collected and stored at −20 °C until further use. Starch was quantified following the modification of Hatterscheid and Willenbrink (1991), for the R-Biopharm quantification kit (10207748035).

#### Semi-high-throughput enzyme activity profiling

A sequential protein extraction was applied following Jammer et al. (2015) and Fimognari et al. (2020), the initial ratio of sample weight to extraction buffer being adjusted to 0.2 g per 1.5 ml. The 1^st^ extraction was performed under low ionic strength conditions in 40mM Tris-HCl pH 7.6, 1mM EDTA, 0.1mM PMSF, 1mM benzamidine, 14mM β-mercaptoethanol, 24 μM NADP and 0.1% PVPP. The resulting pellet was washed three times with distilled water and ionically bound proteins extracted in 1ml of 40mM Tris-HCl pH 7.6, 3mM MgCl_2_, 1mM EDTA and 1M NaCl. Both protein fractions were dialysed overnight against 20 mM potassium phosphate buffer pH 7.4, aliquoted and kept at −20°C until further use. According to Jammer et al. (2015), and in 96-well microtiter plates, the activity of carbon metabolism was estimated using 2-6 ml dialysed aliquots (depending on the enzyme and cellular extract). The activities of Fructose 1,6-bisphosphate aldolase (ALD, EC 4.1.2.13), Glucose-6-phosphate dehydrogenase (G6PDH, EC 1.1.1.49), Phosphoglucoisomerase (PGI, EC 5.3.1.9), Phosphoglucomutase (PGM, EC 5.4.2.2), UDP-glucose pyrophosphorylase (UGPase, EC 2.7.7.9), ADP-glucose pyrophosphorylase (AGPase, EC 2.7.7.27), Sucrose synthase (SuSy, EC 2.4.1.13) and the three isoforms of β-fructofuranosidase also known as Invertase (EC 3.2.1.26; cell wall, cwINV; vacuolar, vacINV; cytosolic, cINV) were quantified and expressed as nkat.g^-1^ seed.

#### Semi-high-throughput antioxidant profiling and malondialdehyde (MDA) determination

To assess the antioxidant capacity, a single extraction protocol was used (Amby et al., 2025). Briefly, 250 µl of 100% methanol was added to 100 mg grounded seeds and the samples were incubated for 1h on a rotating shaker at 4°C. After centrifugation (1600*g*, 5min, 4°C), the supernatant was collected, and the procedure repeated. Pooled supernatants were stored at −20°C until further use. Antioxidant capacity was estimated in 96-well microtiter plates using 2.5-10 µl aliquots and suitable standard curves with the reference standard compounds: gallic acid for Folin-Ciocalteau, 6-hydrozy-2,5,7,8-tetramethyl chroman-2-carboxylic acid for trolox equivalent antioxidant capacity (TEAC), cyanidin-3-glucoside equivalents for total anthocyanins, catechin for total flavanols.

MDA content, a proxy of lipid peroxidation, was determined in a microplate assay adapted from Heath and Packer (1968). Briefly, 1 ml of 1% TCA (trichloroacetic acid) was added to 100 mg grounded seeds and the samples were incubated for 10 min on a rotating shaker at 4°C. After centrifugation (1600*g*, 15min, 4°C), 200 µl of the supernatant and 800 µl of 0.5% of TBA (thiobarbituric acid) in 20% TCA were incubated at 95°C for 30 min followed by 10 min incubation on ice. After centrifugation (1600*g*, 10min, 4°C), 200 µl of the supernatant were added to a microplate and Absorbance recorded. Following non-specific background subtraction (Abs532-Abs600), MDA-TBA complex was estimated using the extinction coefficient of 155 mM^-1^ cm^-1^.

#### Seed nutrient density score

The nutrient density score, NDS_1_, represent a modification of the score proposed by Fulgoni 3^rd^ et al. (2009). It makes use of all the essential minerals quantified in our dataset, as well as introducing antioxidant status as a parameter. Within each parameter and genotype, each value is z-score transformed according to the formula z = (x - x^-^) / s, where **x^-^** is the sample mean, and **s** is the sample standard deviation. For protein estimation, a 5.4 nitrogen-to-protein conversion factor was applied as this average conversion factor is considered to better reflect legume seed protein content (Mariotti et al. 2008). Starch, considered a negative to neutral nutrient, was weighted to 0.5. As legume seeds are rich in health promoting antioxidants (e.g. Ajay et al, 2024), the TEAC/MDA ratio, calculated using the same unit, and weighted to 0.05 was included.

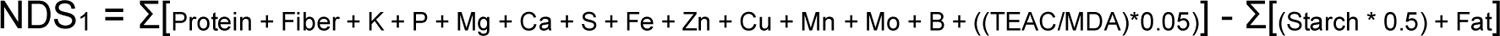

To compare the nutrient density with information available in the literature, for which only a subset of these parameters are available, NDS_2_ was calculated as follows:

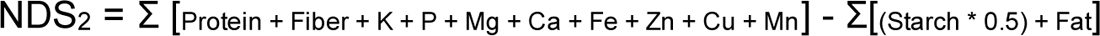

For this, we made use of information available at FoodData Central database (https://fdc.nal.usda.gov/, April 2025) under the NDB Number 100311 and FDC Sample ID 2644311, 2644313, 2644315, 2644317, 2644319, 2644321, 2644323, 2644325. Data from Yegrem (2021), Devi et al. (2023) and Brun et al. (2024) was also used.

### Statistical analysis

All analyses were conducted using R Statistical Software (version 4.2.0; R Core Team 2021). Principal component analysis (PCA) was performed with the Ade4TkGUI package (Thioulouse and Dray, 2007). The Pearson’s product-moment correlation coefficient was calculated using the cor.test function from the stats package. One-way ANOVA and Fisher’s least significant difference (LSD) post hoc tests were carried out with the agricolae package (Mendiburu and Yaseen, 2020).

## Results

### Chickpea reproductive development under controlled conditions

Exposure of chickpea plants to post-flowering elevated temperatures at the Phenolab resulted in reproduction cycle completion within 30-35 days, regardless of watering regime or genotype. Under the GH, it took additional 30 days to complete this cycle. When compared with the GH assay, it was observed that under the high temperature conditions both genotypes did not engage in a water saving strategy (Figure 2). At the Phenolab and following the imposition of the high-temperature cycle (from d08 onwards), leaf temperature increased in both genotypes and watering regimes (Supplemental Figure S1A-H). Figure 2 also illustrates that, regardless, growing conditions (Phenolab or GH) and genotype, water consumption began to decline from d19 onwards. Thus, although under GH conditions plants remained green without visible signs of leaf senescence, the two genotypes had reached their peak capacity to extract water from the substrate, decreasing water spending from now on.

**Figure 2.**
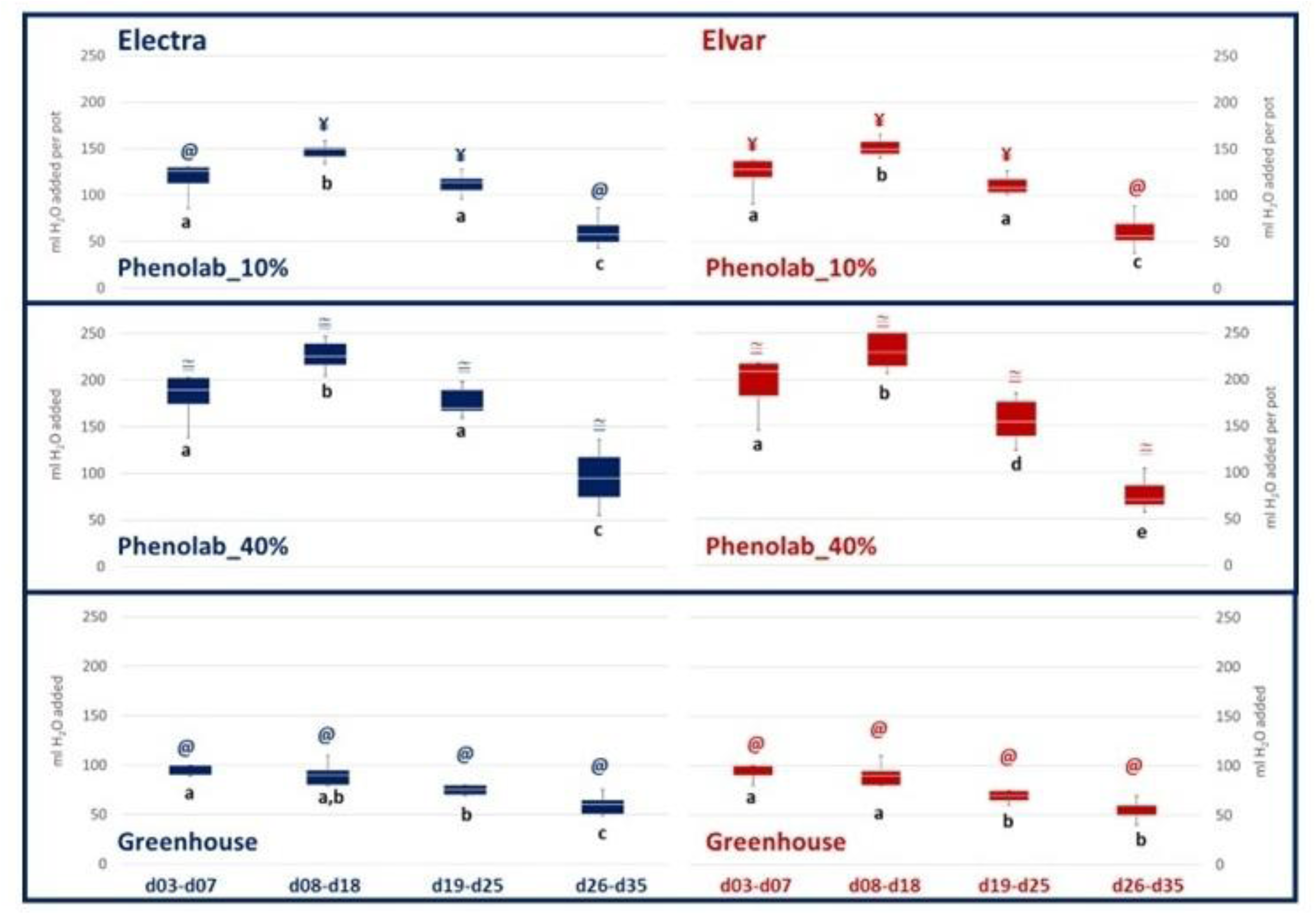
Post-flowering water spending for Electra and Elvar genotypes under different irrigation and temperature conditions. The genotypes were grown under greenhouse conditions with 40% full irrigation and at the Phenolab under high temperatures with either 40% (Phenolab_40%) or 10% (Phenolab_10%) full irrigation. The greenhouse growth cycle concluded at day 65, while the Phenolab cycle was concluded 35 days after flowering. One-way ANOVA followed by Fisher’s least significant difference (LSD) post hoc tests were used to determine significant differences at p < 0.05. Lowercase letters below the mean values compare genotypes under the same treatment and period, while symbols above the mean values compare treatments within the same genotype and period.

The multispectral data gathered through the reproductive development under high temperatures at the Phenolab was used to calculate several vegetation indices (Supplemental Table S4). A PCA analysis (Figure 3) showed that six indices closely clustered together: NVDI (normalised difference vegetation index), NRDE (normalised difference red-edge vegetation index), GNDVI (green normalized difference vegetation index), EVI (enhanced vegetation index), DSWI-4 (disease-water stress index 4) and RVI (ratio vegetation index). Along PC1 and PC2 (67% and 15% of the variation explained, respectively), the multispectral data also evidenced clusters that corresponded to distinct phases of water consumption, as illustrated in Figure 2. The first phase (d01-d06) corresponds to the acclimation periods. The second phase (d07-d18) corresponds to the period when plants used the highest amount of water. In the third phase (d19-d25), water consumption began to decline, with noticeable differences between Electra 40% and Elvar 40% emerging in the later days. Finally, during the last phase (days 27–35), plants under the 40% water regime exhibited similar water consumption regardless of temperature. Four vegetation indexes provided the clearest differences between genotypes: ARI (anthocyanin reflectance index), PRI (photochemical reflectance index), WBI4 (water band index 4) and iRECI (inverted red-edge chlorophyll index; Figure 4, Supplemental Figure S2). ARI consistently distinguished the genotypes along the reproductive development (d01-d35), while PRI and WBI4 showed a marked effect of increasing the temperature (before and after d07). PRI and WBI4 also exhibited genotype related differences, more evident from d02-d25 and from d16-d29, respectively. iRECI revealed genotype differences later in the growing cycle, particularly from d26-d35.

**Figure 3.**
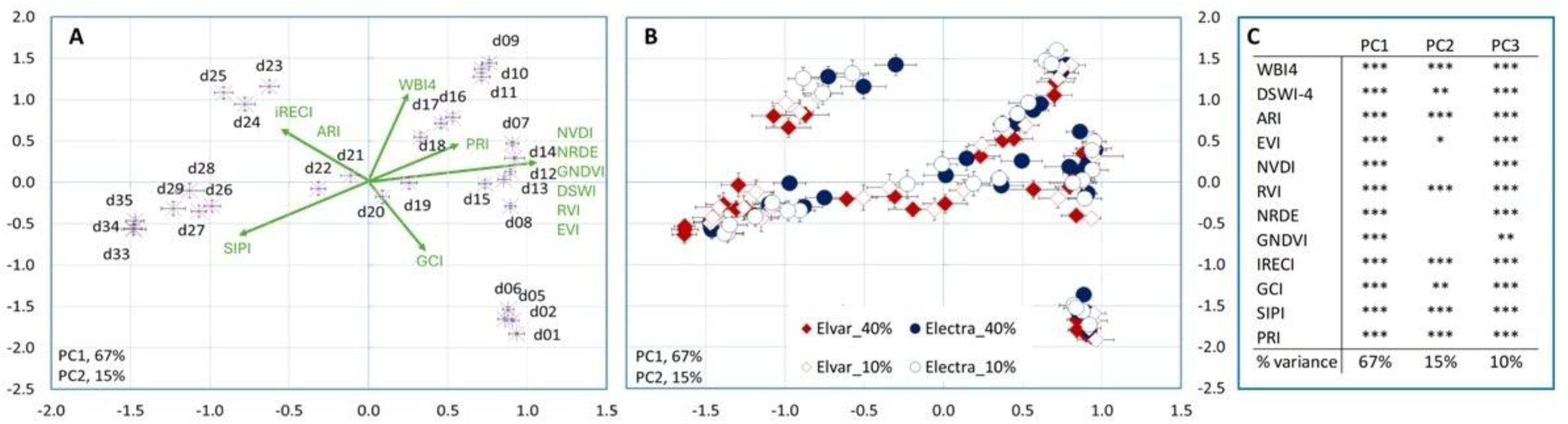
Principal components analysis (PCA) making use of 12 vegetation indices according to the number of days in the Phenolab (**A**) or according to the number of days, genotype and treatment (**B**). In (**C**), the Pearson’s product moment correlation coefficient for each variable (*p < 0.05, **p < 0.01; ***p < 0.001). In (**A**), the indices significantly contributing to the separation along PC1 and PC2 are displayed in green. Multispectral data used as input for the calculation of the vegetation indices is available as Supplemental Table S4).

**Figure 4.**
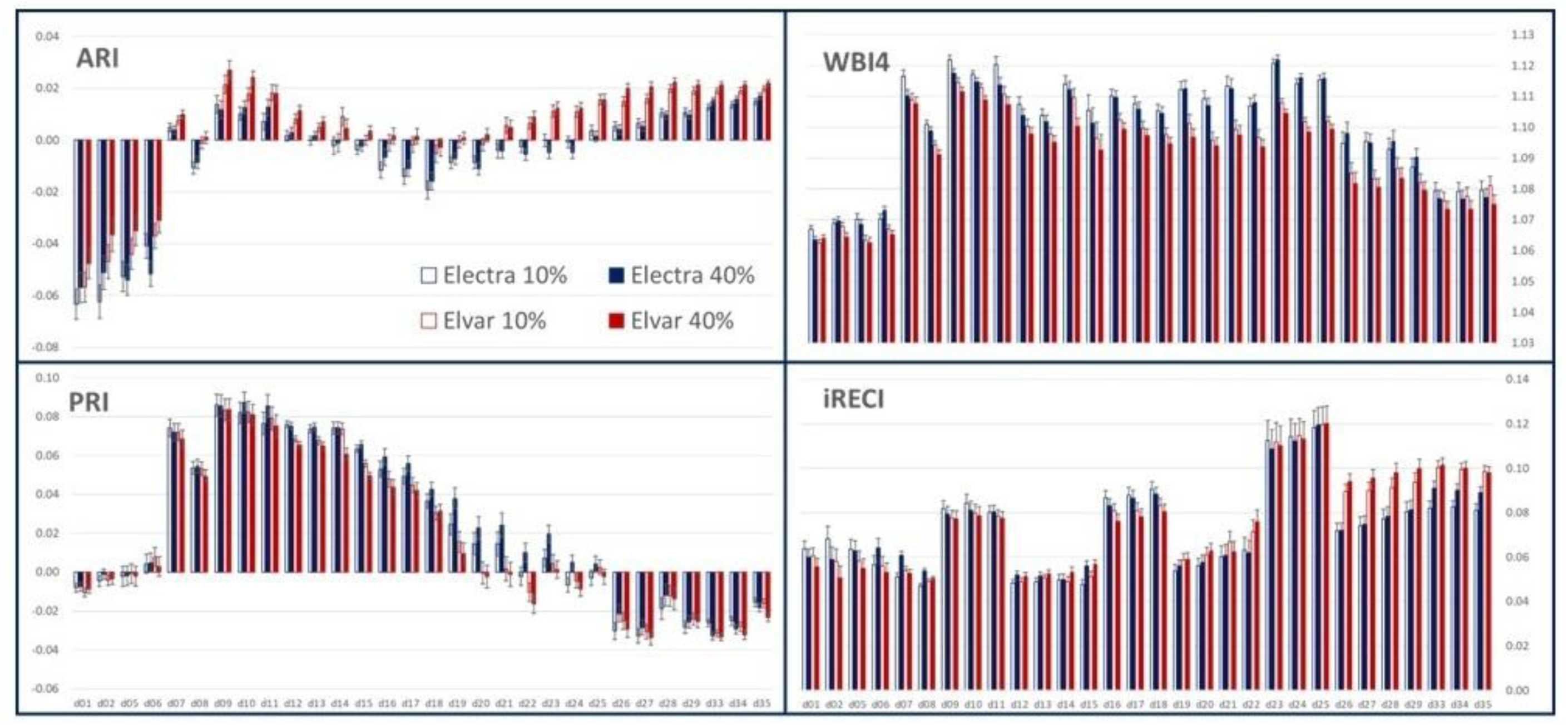
Selected vegetation indices for Elvar and Electra from flowering (d01) to cycle completion (d35) under post-flowering high temperatures and 40% full irrigation (Phenolab_40%) or 10% full irrigation (Phenolab_10%). Full data is available as Supplementary Table S4. Additional indices are available as Supplementary Figure S2.

### Chickpea seed phenotype, yield and quality

For both genotypes, the highest 100-seed weight was observed under GH conditions, the exposure of chickpea plants to post-flowering high temperatures negatively impacted not only the weight of 100 g seeds but also the number of dry seeds per plant (Table 1). However, a further effect of the watering regime was only evidenced on Elvar, the lowest 100-seed weight were obtained from plants under the 40% watering regime.

**Table 1.**
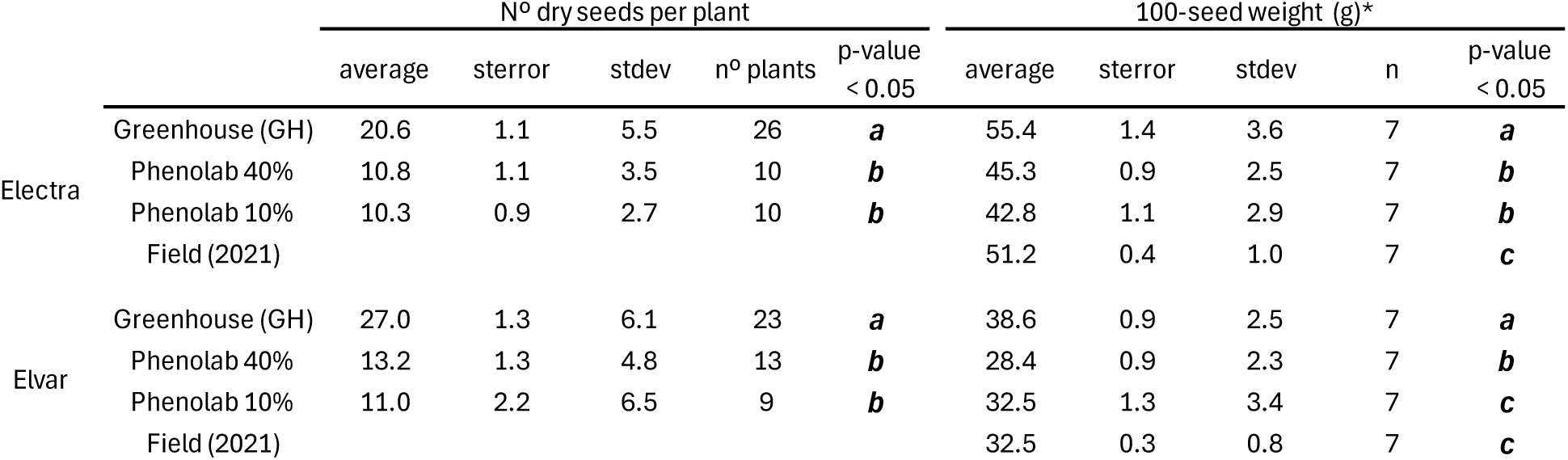
Number of dry seeds per plant and 100-seed weight (g) for two chickpea genotypes (Electra and Elvar) grown under different environmental conditions: greenhouse (GH), Phenolab with 40% and 10% irrigation, and field conditions (2021). Values are average with standard error (sterror) and standard deviation (stdev). The number of plants (n° plants) or replicates (n) used for each condition is also indicated. One-way ANOVA (GxT, p<0.001) was followed by Fisher’s least significant difference (LSD) post hoc tests. Different letters within each genotype indicate statistically significant differences among treatments (p < 0.05).

|  |  | N° dry seeds per plant |  |  |  |  | 100-seed weight (g)* |  |  |  |  |
| --- | --- | --- | --- | --- | --- | --- | --- | --- | --- | --- | --- |
|  |  | average | sterror | stdev | n° plants | p-value<br>< 0.05 | average | sterror | stdev | n | p-value<br>< 0.05 |
| Electra | Greenhouse (GH) | 20.6 | 1.1 | 5.5 | 26 | **a** | 55.4 | 1.4 | 3.6 | 7 | **a** |
|  | Phenolab 40% | 10.8 | 1.1 | 3.5 | 10 | **b** | 45.3 | 0.9 | 2.5 | 7 | **b** |
|  | Phenolab 10% | 10.3 | 0.9 | 2.7 | 10 | **b** | 42.8 | 1.1 | 2.9 | 7 | **b** |
|  | Field (2021) |  |  |  |  |  | 51.2 | 0.4 | 1.0 | 7 | **c** |
| Elvar | Greenhouse (GH) | 27.0 | 1.3 | 6.1 | 23 | **a** | 38.6 | 0.9 | 2.5 | 7 | **a** |
|  | Phenolab 40% | 13.2 | 1.3 | 4.8 | 13 | **b** | 28.4 | 0.9 | 2.3 | 7 | **b** |
|  | Phenolab 10% | 11.0 | 2.2 | 6.5 | 9 | **b** | 32.5 | 1.3 | 3.4 | 7 | **c** |
|  | Field (2021) |  |  |  |  |  | 32.5 | 0.3 | 0.8 | 7 | **c** |

Although the post-flowering high temperatures did not affect seed germinative potential, it affected the seed colour, size, composition and biochemistry (Figure 5, Supplementary Tables S5.1 to S5.4). As clearly illustrated by Figure 5, and when produced under the same growing conditions, Electra seeds are bigger than those from Elvar. It was also observed that seeds produced under field conditions were lighter than the ones obtained under controlled conditions, the most striking colour alteration being register for Elvar seeds obtained under the combination of post-flowering high temperatures and 10% watering regime. In addition, seeds obtained under post-flowering high temperatures exhibited a wrinkled phenotype, particularly for the Electra genotype.

**Figure 5.**
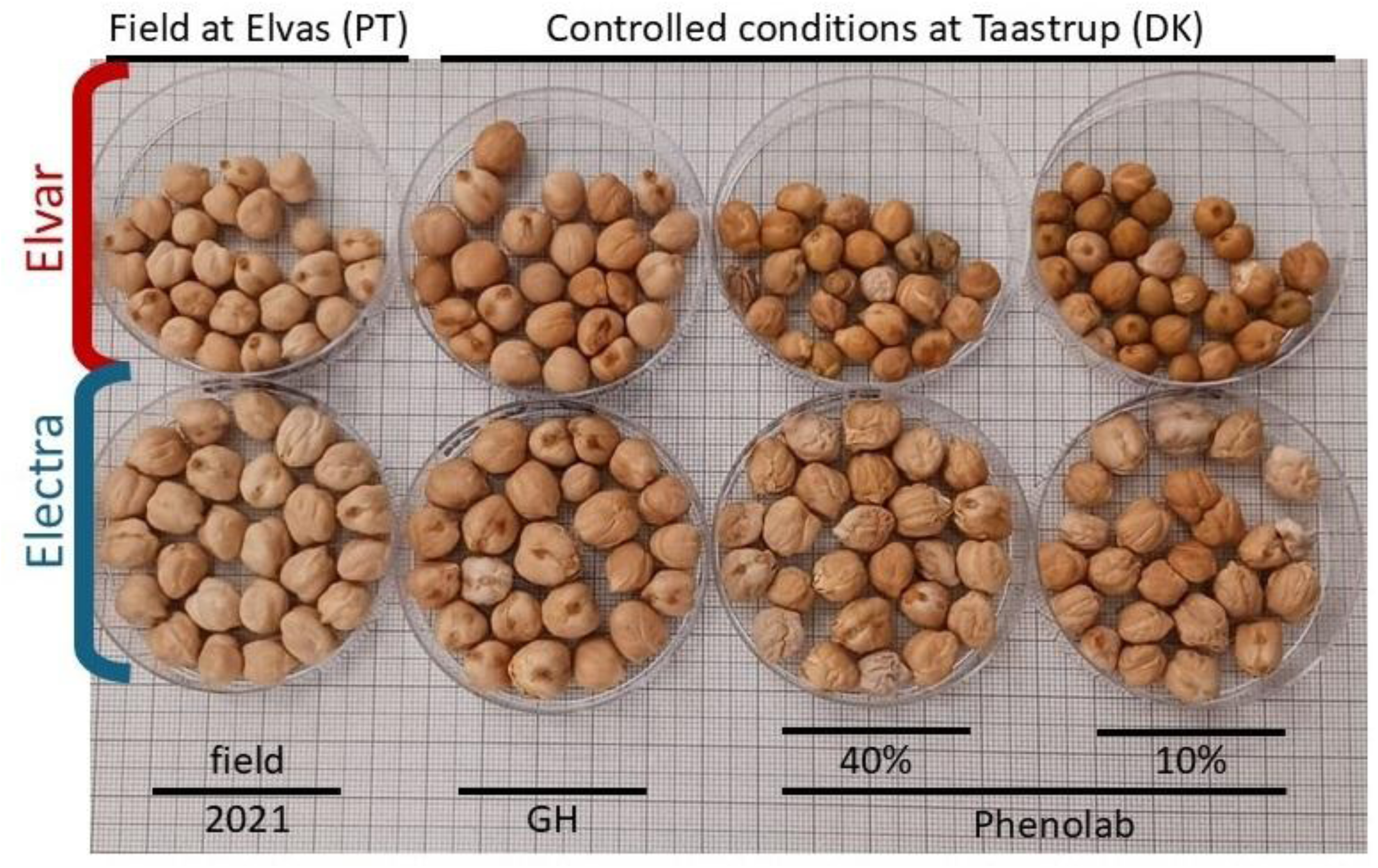
Phenotypes of dry and mature seeds of Electra and Elvar genotypes. GH – seeds obtained from plants maintained in the greenhouse; 40% and 10% – seeds obtained from plants submitted to post flowering heat treatment at the Phenolab under a 40% or 10% full watering regime, respectively; 2021 – seeds obtained from plants grown under field conditions at Elvas in the 2021 growing season. For all treatments, parental seeds were obtained from plants grown under field conditions at Elvas in the 2020 growing season.

PCA analysis (Figure 6) of seed composition and biochemistry reveals that seeds exposed to post-flowering high temperatures cluster together based on their genotype. Along PC1 (31% of the variation explained), seeds obtained under post-flowering high temperatures are separated from the ones obtained from plants grown in field and GH conditions, while along PC2 (21%) seeds were separated according to their genotype. However, our analysis identifies two main groups of variables: seed carbon metabolism enzyme activities that mainly reflect seed genotype (Supplemental Figure S3); seed composition that primarily reflects the plants’ growing conditions, regardless of genotype, and with a clear influence of the watering regime under post-flowering high temperatures (Supplemental Figure S4).

**Figure 6.**
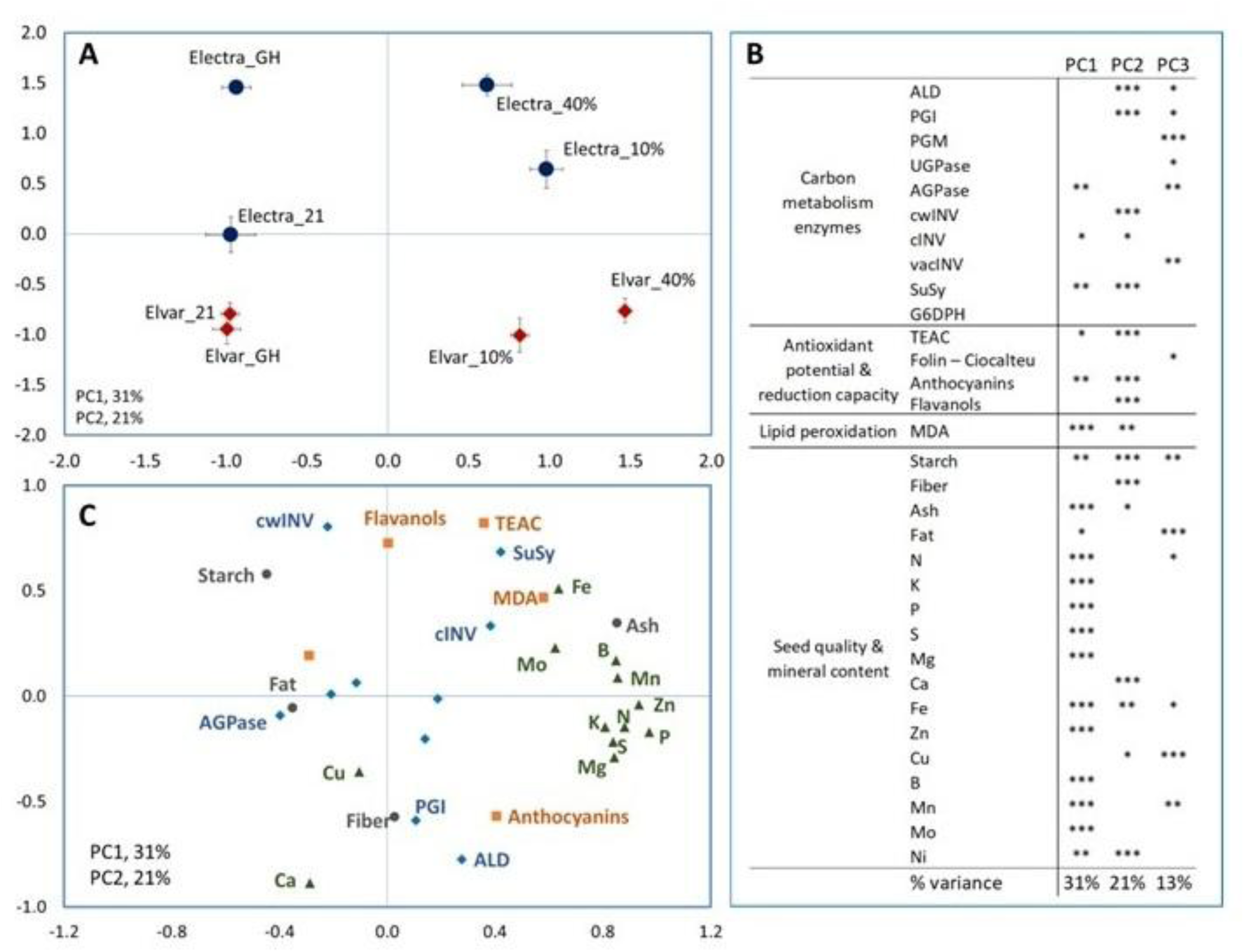
Principal components analysis (PCA) and making use of biochemical and composition data from completely dry and mature seeds of Electra and Elvar genotypes (**A**) and the Pearson’s product moment correlation coefficient for each variable (*p < 0.05, **p < 0.01; ***p < 0.001; (**B**). Variables significantly contributing to the separation along PC1 and PC2 are presented in (**C**). GH, seeds obtained from plants maintained in the greenhouse; 40% and 10%, seeds obtained from plants submitted to post flowering heat treatment at the Phenolab under a 40% or 10% full watering regime, respectively; 21, seeds obtained from plants grown under field conditions at Elvas in the 2021 growing season. The parental seeds were the same for all treatments and were seeds obtained from plants grown under field conditions at Elvas in the 2020 growing season. Data used as input for the calculation of the vegetation indices is available as Supplemental Tables S5.1 to S5.4.

Post-flowering high temperatures had a positive effect on seed nutrient density (Figure 7), with water availability further modulating this response. In our dataset, genotypic differences were evident under all conditions except for the 40% watering regime and high post-flowering temperatures. The typical trade-off between protein (organic nitrogen content, estimated by the Kjeldahl method) and starch content or seed weight was observed (Table 1, Supplementary Table S5.1). Additionally, and although the magnitude of these effects varied, the impact of post-flowering high temperatures on seed composition (N, starch, fiber, fat, ash, and minerals) generally followed the same trend, either increasing or decreasing in both genotypes (Supplementary Table S5.1 and S5.2). Seeds from both genotypes exhibited relatively higher mineral content when developed under post-flowering high temperatures. In Elvar, mineral accumulation was larger under the 40% watering regime (Ca was the only mineral showing a significant decreased comped to GH), whereas in Electra, it was higher under the 10% watering regime, due to all but Ca macronutrient’s increase. Micronutrient content was water responsive in both genotypes, generally being higher under the 40% watering regime, except for Cu and B in Elvar.

**Figure 7.**
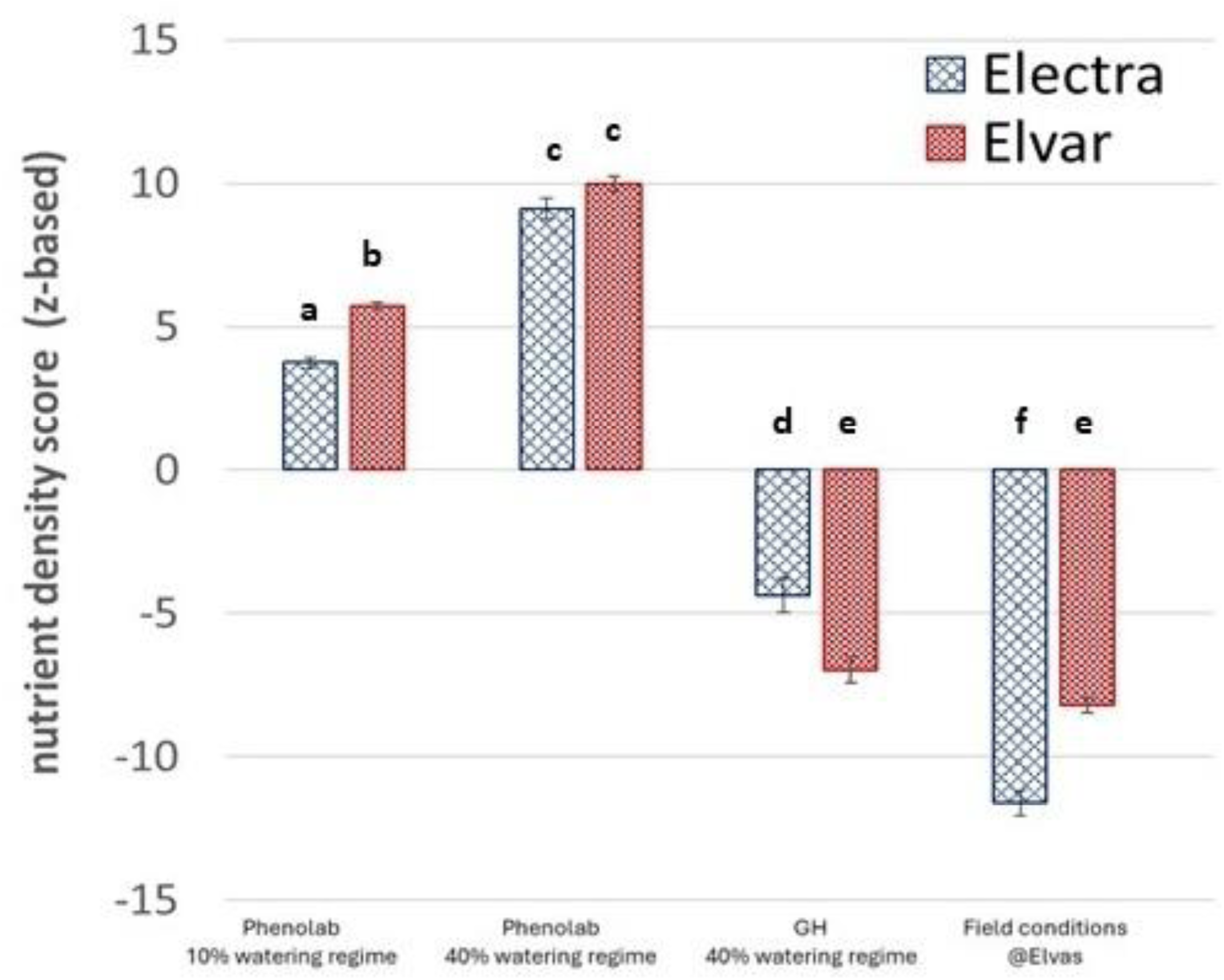
Nutrient density score NDS_1_ for completely dry and mature seeds of Electra and Elvar. NDS_1_ was calculated using the z-scores of the several parameters. One-way ANOVA followed by Fisher’s least significant difference (LSD) post hoc tests were used to determine significant differences at p < 0.001. GH, seeds obtained from plants maintained in the greenhouse; 40% and 10%, seeds obtained from plants submitted to post flowering heat treatment at the Phenolab under a 40% or 10% full watering regime, respectively; Field, seeds obtained from plants grown under field conditions at Elvas in the 2021 growing season. The parental seeds were the same for all treatments and were seeds obtained from plants grown under field conditions at Elvas in the 2020 growing season. Data used as input is available as Supplemental Tables S5.1 and S5.2.

As previously referred the seed carbon metabolism enzyme activities mainly reflect genotypes, that clearly separated along PC1, explaining 28% of the variation (Supplemental Figure S3). Enzymes that most contributed for the differentiation of the genotype were: cell wall invertases (cwINV), cytosolic invertases (cINV) and sucrose synthase (SuSy), with higher activity in Electra; and fructose 1,6-bisphosphate aldolase (ALD) and phosphoglucoisomerase (PGI), with relatively higher activity in Elvar (Supplemental Figure S3; Supplementary Table S5.4). Considering the INV isoforms, cwINV was found to be the most active isoform in seeds in both genotypes. Enzymatic activities profile also reveals that, under the tested conditions, seed variability was higher in Electra than in Elvar, particularly due to Electra seeds obtained under field conditions.

In addition to carbon enzyme activities, the antioxidant capacity and MDA content was also assessed (Supplementary Table S5.3). A proxy of membrane lipid peroxidation, MDA content was found to be increased in seeds developed under post-flowering high temperatures (> 2-fold in Elvar; >1.5-fold in Electra), irrespectively of the watering regime. The antioxidant capacity was estimated using several biochemical methods. While each method provided a distinct response profile, all consistently highlighted the influence of genotype and growing conditions on the seed antioxidant capacity. The antioxidant potential (calculated as the sum of trolox, gallic acid, cyanidin-3-glucoside and catechin equivalents) was not affected by the post-flowering high temperatures in seeds from Elvar but it decreased in seeds from Electra obtained under the 10% watering regime. This effect can be related with a decrease in antioxidant capacity as estimated as gallic acid equivalents (the Folin-Ciocalteau method) as it represented for > 80% of the antioxidant potential (Supplementary Table S5.3). When comparing to seeds developed under GH conditions, and for both genotypes, the post-flowering high temperatures lead to higher trolox equivalents under the 40% watering regime. The flavanols content, calculated as catechin equivalents, were found to be genotype and watering regime dependent, its content being increased in seeds from Elvar obtained under the 10% watering regime. In Electra, anthocyanins (calculated as cyanidin-3-glucoside equivalents) were not detected in seeds obtained under the GH conditions, and for the remaining treatments the content was found to be ten times lower than the flavanols. This profile was distinct in Elvar, that exhibited a relatively higher anthocyanin content. Although seed colour was affected by growing the plants under high temperatures (Figure 5), the low anthocyanin content and response profile does not allow to fully explain the observed colour changes.

## Discussion

Understanding how chickpea plants respond to post-flowering high temperatures and limited water availability is crucial for developing strategies to ensure sustainable yield and seed quality under increasingly challenging climatic conditions. The duration of the reproductive phase is a key factor in determining seed yield (Devasirvatham et al., 2015; Maity et al. 2023; Sita et al., 2017). Screening genetic diversity for tolerant genotypes is necessary, as even those well adapted to hot and dry conditions, such as Elvar and Electra, may fail to secure yield. Consistent with previous findings (e.g. Awasthi et al. 2014, 2017; Benali et al. 2023), our data show that yield losses can exceed 50% when using number of seeds per plant and the weight of 100 seeds as yield proxies. Although chickpea grain yield responds positively to soil water availability (Silva et al. 2014), it is also indicated that reductions in the reproductive phase duration caused by high post-flowering temperatures may not be fully mitigated by irrigation. Although chickpea plants typically continue growing and using water throughout the reproductive phase due to their indeterminate growth habit, both Elvar and Electra showed a sharp and simultaneous decline in maximum water use. This decline occurred independently of temperature or water availability, suggesting a time-limited physiological or genetic constraint on the maximum water uptake during the reproductive phase. Previous studies shown that chickpea drought tolerant genotypes have a conservative water strategy, not using all the water available and promoting sustainable water use into the reproductive stage (e.g. Zaman-Allah et al. 2011).

In Electra, the weight of 100 seeds followed a more understandable pattern, with the greatest reduction in seed weight occurring under combined stresses (post-flowering high temperature and limited water availability), while single stress factors resulted in intermediate negative effects. In contrast, Elvar exhibited a more complex response. Both the maximum and minimum seed weights observed under the 40% watering regime, suggesting a more intricate relationship with water availability. The better performance of Elvar under the most extreme condition tested suggests that the genotype adaptation potential may be further explored. However, seed phenotype (colour and size) may hamper its acceptance.

While the negative impact of high temperatures and water availability on the duration of the chickpea reproductive phase have been described as early 1980’s (e.g. Summerfield et al. 1984), their nutritional implications remain far less explored. In the literature, protein and starch content in response to environmental conditions are the more frequently reported seed nutritional parameters. Consistent with our findings, previous studies have reported protein-to-starch trade-offs in chickpea (Frimpong et al. 2009; Wang et al. 2017). These studies identified a positive correlation between starch content and seed yield, and a negative correlation between starch and protein content, with significant G × E interactions that explained more variance than genotype effects alone. Domergue et al. (2019), reviewing the high temperatures impact on seed quality, report that such conditions lead to increasing allocation of assimilates protein and away from non-structural carbohydrates. However, not all studies observed such trade-offs, instead reporting a concurrent reduction in yield, starch, and protein content under stressful conditions (e.g., Devi et al. 2023; Awasthi et al. 2024).

The seed nutrient density strongly reflects genotype growing conditions, the post-flowering high temperatures leading to a higher nutrient density in our data set. The comparison with other values in the literature was difficulted due to data availability and consistency in the reported parameters. The difficulty in data access and re-use is consider one major obstacle and several efforts are on-going to make data FAIR compliant, i.e. Findable, Accessible, Interoperable, and Reusable (e.g. Papoutsoglou et al. 2020, Top et al. 2022). Nevertheless, for chickpea and based on the average values reported by Devi et al. (2023) for protein, starch, fat, fiber, Ca, P, Fe, a similar trend seems to be observed for the effect of post-flowering high temperatures. Literature comparisons with available data also reinforce the significant influence of G × E interactions on seed nutritional potential (Supplemental Figure S5). Genotypes selected under the climate conditions of the Iberian Peninsula (Spain and Portugal) were found to exhibit comparable nutrient density levels, but notably different from those reported by Yegrem (2021) and Devi et al. (2023) for chickpea genotypes grown in Ethiopia and India, respectively. When applying the same analytical approach to data from the Food Central Database (for seeds produced in Virginia, USA, in 2023), nutrient densities were found to be lower than those from the Iberian Peninsula, but higher than those from India and Ethiopia. Significant advances can be made by addressing both the technical and structural barriers to data availability and integration as researchers can make meaningful progress in identifying and selecting genotypes for nutritional quality and technological traits under changing climates.

The shortening in the seed filing period caused by post-flowering high temperatures is considered a major cause for yield reduction as reviewed in Sita et al. (2017). At the biochemical level, that is attributed to lower starch accumulation since starch is a major contributor for seeds dry weight. In our dataset, a direct the relationship between these two parameters is not evidenced. This was already contextualised by Kumar et al. (2023) on their revision on post-flowering high temperatures impacts on cereals seeds. Seed starch accumulation reflects the seed sink strength in the context of plant photosynthetic capacity and source distribution. In addition, sucrose imported into the seed will be channelled for distinct blocks (e.g. starch or protein synthesis). Seed sink strength is related with the ability to maintain the sucrose gradient favourable to sucrose import into the seeds. According to Kumar el al. (2023), genotypes with increased seed vacuolar invertase activity can secure starch synthesis under stress. Devi et al. (2023) and Awasthi et al. (2024) also evidenced reduced activity of this enzyme (along with reduced activity of SuSy) in chickpea seeds developed under elevated post-flowering temperatures as a cause for lower yield and starch content. In our dataset, however, despite the decrease in starch content, distinct enzymatic profiles observed between the two genotypes analysed. Most likely, multifactorial regulations of enzyme activities that modulate sink strength and reserve accumulation are to be considered. According to the speciés ideotype, monitor carbon assimilation potential and its distribution to pods and seeds has the potential to disclose metabolic patterns favourable to seed set and reserve accumulation. This divergence may reflect genotype-specific regulatory mechanisms, potentially linked to the genetic backgrounds and selection histories of the genotypes. The large difference in seed ALD activity between the Electra and Elvar can highlight such differences. In the literature, high ALD activity has been associated with seed readiness for germination (Krüger and Schanarrenberger 1985), as it supports the rapid resumption of glycolysis once favourable conditions return. Additionally, ALD activity may contribute to membrane protection during the desiccation process, providing sugar skeletons and energy for osmoprotectants accumulation. In our dataset, and under controlled conditions, higher ALD activity co-occurred with lower malondialdehyde (MDA) levels in Elvar seeds compared to Electra seeds harvested under the same conditions, suggesting a possible link between metabolic priming and oxidative stress mitigation. A more detailed analysis of the evaluated genotypes, including their geographical origin and selection ideotypes, could provide valuable insights into these differences and inform their potential utility in future breeding programs. While not sufficient to adequately reflect seed physiological quality, where metabolic fluxes, rather than single metabolic features, are relevant (e.g. Domergue et al. 2019), enzymatic activity profiles can still serve as valuable indicators of underlying genotype - specific metabolic regulations, e.g. in leaves (Correia et al. 2022, Rasool et al. 2024). To the best of our knowledge, our study is the first to demonstrate the utility of seed carbon metabolism enzymes in discriminating genotypes.

## Conclusions

The reproductive development of two elite chickpea genotypes, Electra and Elvar, adapted to the dry conditions of southern Portugal and known for their high production capacity, was evaluated under post-flowering high temperatures and differential watering regimes. Overall, our findings demonstrate that post-flowering high temperatures significantly impact seed traits, including weight, composition, biochemistry, and enzymatic activity, with genotype and watering regime dependent effects. Electra seeds generally exhibited greater starch and nitrogen responsiveness to high temperatures, highlighting the distinct metabolic profiles of each genotype, reinforcing the role of specific carbon metabolism enzymes in genotype differentiation at the mature seed stage. Reduced assimilate partitioning due to impaired photosynthesis and disruptions in carbon, water, and nutrient transport can weaken sink strength, leading to lower accumulation of essential macronutrients (carbohydrates, lipids) in seeds. However, post-flowering high temperatures may modify metabolic pathways involved in protein synthesis and secondary metabolite production, potentially affecting seed composition and overall nutritional value. In line with existing literature, our findings demonstrate that chickpea yield and nutritional quality are highly sensitive to the combined effects of post-flowering high temperatures and water availability. Furthermore, seed phenotype and composition are crucial indicators of crop performance under stress conditions. The combined effects of high temperatures and altered water regimes can modify seed size, color, biochemical composition and enzyme activities, impacting nutritional quality and economic value. Our approach revealed that, at mature seed level and considering seed composition, genetic differences are overridden by the environment. To ensure stable chickpea production and maintain nutritional value under changing environmental conditions, targeted breeding efforts and adaptive management strategies are essential to pave the way for climate-resilient, nutritionally secure chickpea production.

## Supplementary data

Table S1. Average monthly and annual rainfall (mm) and temperature (°C) in the Elvas region during seed production years.

Table S2. Greenhouse ambient conditions during the growth period of chickpea Electra and Elvar (June–October 2021).

Table S3. Phenolab ambient conditions for chickpea Electra and Elvar under controlled assay (July–October 2021).

Table S4. Multispectral data and vegetation indices for chickpea Electra and Elvar grown at the Phenolab under high-temperature treatment.

Table S5.1. Seed quality and yield-related parameters of Electra genotype under greenhouse, Phenolab, and field conditions.

Table S5.2. Seed quality and yield-related parameters of Elvar genotype under greenhouse, Phenolab, and field conditions.

Table S5.3. Comparative analysis of seed nutrient composition across irrigation and temperature regimes.

Table S5.4. Carbon metabolism enzymatic activity in seeds of Electra and Elvar across environments.

---

**Figure S1.** Leaf temperature dynamics derived from thermal extraction matched with multispectral imaging under a controlled heat increase regime.

**Figure S2.** Vegetation indices of Elvar and Electra across the reproductive cycle under differential irrigation and heat treatments.

**Figure S3.** Principal component analysis of carbon metabolism enzymatic activity in Electra and Elvar seeds, with correlation analysis of contributing variables.

**Figure S4.** Principal component analysis of seed composition data in Electra and Elvar, with correlation analysis of contributing variables.

**Figure S5.** Nutrient density scores (NDS2) of chickpea seeds compared with published data and FoodData Central database values.

## Acknowledgements

At UCPH, Cairo Westergaard and the Taastrup gardening team are acknowledged for their assistance with multispectral analysis and plant growing under controlled conditions. Ana Bagulho (INIAV) is acknowledged for their assistance seed elemental analysis. We acknowledge the Laboratório de Análises/REQUIMTE/LAQV (Carla Rodrigues) for the acquisition of the ICP-OES data. The Bachelor student José Melo (NOVA-FCT) is acknowledged for his contribution to starch analysis. This work was supported by the European Plant Phenotyping Network 2020 (H2020, grant No. 731013), EMPHASIS-GO (HORIZON-INFRA-2021-DEV-02, No. 101079772), and by Portuguese national funds through FCT (UIDP/04378/2020, UIDB/04378/2020, LA/P/0140/2020, UIDB/04129/2020, UIDP/04129/2020, LA/P/0092/2020).

## Author Contributions

CP, LGG and TR: conceptualization; CP, TR and ID: methodology; CP, ECC, MEMF, LGG and TR: formal analysis; CP, ECC and LGG: investigation; CP, CB, ID and TR: resources; CP, ECC and LGG: data curation; CP and ECC: writing - original draft; All authors: writing - review & editing; CP and TR: supervision; CP and TR: funding acquisition

## Abbreviations

AGPase: ADP-glucose pyrophosphorylase
ALD: Fructose 1,6-bisphosphate aldolase
ANOVA: Analysis of Variance
ARI: Anthocyanin Reflectance Index
CCD: Charge-Coupled Device
cINV: Cytosolic Invertase
cwINV: Cell Wall Invertase d01, d02…: Experimental days (day count after start of treatment)
DSWI-4: Disease-Water Stress Index 4
EDTA: Ethylenediaminetetraacetic Acid
EVI: Enhanced Vegetation Index
G6PDH: Glucose-6-phosphate dehydrogenase
GH: Greenhouse
GNDVI: Green Normalized Difference Vegetation Index
ICP-OES: Inductively Coupled Plasma Optical Emission Spectroscopy
iRECI: Inverted Red-Edge Chlorophyll Index
ISO: International Organization for Standardization
K₂O: Potassium Oxide
LSD: Least Significant Difference
MDA: Malondialdehyde
NADP: Nicotinamide Adenine Dinucleotide Phosphate
NDS: Nutrient Density Score
NVDI: Normalized Difference Vegetation Index
PCA: Principal Component Analysis
PGM: Phosphoglucomutase
PGI: Phosphoglucose Isomerase
PMSF: Phenylmethylsulfonyl Fluoride
PRI: Photochemical Reflectance Index
PVPP: Polyvinylpolypyrrolidone
RVI: Ratio Vegetation Index
SuSy: Sucrose Synthase
TBA: Thiobarbituric Acid
TCA: Trichloroacetic Acid
TEAC: Trolox Equivalent Antioxidant Capacity
UGPase: UDP-glucose Pyrophosphorylase
vacINV: Vacuolar Invertase
WBI4: Water Band Index 4

